# Microclimate buffering varies across forest types during an extreme heat event

**DOI:** 10.1101/2023.10.02.560390

**Authors:** Aji John, Kavya Pradhan, Michael J. Case, Ailene K. Ettinger, Janneke Hille Ris Lambers

## Abstract

Increasing temperatures and extreme heat episodes have become more common with climate change. While forests are known to be buffered from increasing temperatures compared to non-forested areas, whether this buffering is maintained under extreme temperature events, how such events influence forests, and how forest organisms respond to extreme heat is relatively unknown. Here we assess the effects of an extreme heat event (the Pacific Northwest (PNW) heatdome in June 2021) on forest microclimates, forests, and the organisms living within them. We first asked how the PNW heatdome affected microclimates in forests with differing canopy cover (including non-forests) and found that the buffering capacity of forests is greater under denser canopies, even under extreme heat events. We then combined this information with organismal temperature tolerance curves for 12 relevant species and found that canopy buffering can minimize the negative impacts of even extreme heat events on understory organisms, with greater canopy density providing greater microclimate moderation. Finally, we analyzed seasonal NDVI trends in recent years, and found signs of canopy stress following the extreme 2021 heat event. In all, this suggests that although forest canopies may buffer the negative effects of extreme heat events on understory organisms, a greater frequency of extreme heat events may threaten this capacity by damaging forest canopies.

## Introduction

Anthropogenic climate change has had strong impacts on forests due to increasing temperatures (Case et al. 2021, De Frenne et al. 2021) as well as altered precipitation regimes (Knapp et al. 2015). For example, rising tree mortality rates have been linked to increases in regional temperatures and climate-driven changes in water availability both regionally and globally (Van Mantgem et al. 2009, Allen et al. 2010, Perkins-Kirkpatrick and Lewis 2020), and treeline has generally moved upward in elevation in response to longer growing seasons associated with warming (Harsch et al. 2009). The consequences of long-term changes in average climates on forests and organisms have received a lot of attention and this rich literature has provided a lot of insight into the longer-term impacts of climate change. However, responses to more intense but shorter duration extreme weather events (like heatwaves and heatdomes (White et al. 2023)) have more unknowns as there are fewer studies that have examined these impacts in a variety of locations. Heatdomes and heatwaves, periods of extremely hot weather, are particularly concerning given the increase in their frequency in recent years and the potential negative impacts on ecosystems worldwide.

How might forest dwelling organisms respond to extreme climate events? Forests are known to provide understory organisms reprieve from warm temperatures with canopy cover buffering the understory from the broader macroclimate (De Frenne et al. 2019) but whether forests maintain their buffering capacity during extreme heat events is an open question. A major reason for the buffering capacity of forests is their dense canopy cover (Davis et al. 2018a, Zellweger et al. 2019, Gril et al. 2023), although topography can also play a role (Greiser et al. 2018, Zellweger et al. 2019). The forest canopy intercepts incoming solar radiation, allowing only diffuse radiation to reach the understory (Lieffers et al. 1999, Drever and Lertzman 2003), and additionally evaporatively cools (through transpiration of canopy leaves) the understory (Oke 1976, Chen et al. 1999), thus keeping areas with canopy cooler and less exposed to temperature fluctuations than areas without. Previous studies have found lower maximum (Renaud and Rebetez 2009, Davis et al. 2018a) and higher minimum temperatures (De Frenne et al. 2019) in areas with canopy cover when compared to areas without canopy cover. Similarly, forest understories are known to have lower diel variation in temperature (max daily – min daily temperatures) due to increased long-wave:shortwave radiation (absorption of shortwave radiation for photosynthesis), potentially leading to more stable climatic conditions (Scheffers et al. 2014). There is also evidence that these effects are particularly pronounced when regional conditions are warm (Renaud and Rebetez 2009, Ashcroft et al. 2012), suggesting that forests may play a similar buffering role during extreme weather events. If so, perhaps forests can buffer understory organisms from extreme heat events through the moderating effects of their canopies. Moreover, if forest canopies can persist and maintain their buffering capacity through these increasingly common extreme, but short-duration climatic events, then it is possible that extreme heat events would not impose an additional burden on organisms beyond that caused by increasing mean temperatures.

Recent studies have also found that forest characteristics matter for the level of understory microclimate buffering. As canopy characteristics vary across forests, and differences in tree density can alter the amount of solar radiation reaching the forest floor. These differences have been quantified in European deciduous forests showing the cooling effect of more shade during summer (Zellweger et al. 2019). In addition to this, differences in stand characteristics, including shade casting ability, can impact the relationship between macroclimate and microclimate under forest canopies (Gril et al. 2023). However, the relative importance of variable canopy cover on microclimate buffering during extreme thermal events is unknown. Forest buffering capacity has been linked to long term persistence of biodiversity (Betts et al. 2018, Zellweger et al. 2020), but it is uncertain whether this buffering is maintained enough to prevent damage to understory organisms during heatwaves and heatdomes. Extreme heat events can impact understory organisms through effects on physiology (Reyer et al. 2013), particularly if temperatures rise beyond the maximum thermal tolerances of organisms. Such physiological impacts are pointed to as causes for heatwave associated mortality observed in birds (McKechnie and Wolf 2010) and small mammals (Ratnayake et al. 2019, Czenze et al. 2022). Similarly, few studies have directly assessed the impact of extreme heat events on forests (Margalef-Marrase et al. 2020, Yi et al. 2022). Those that have found divergent results. For example, one study found that higher than normal temperatures associated with heat waves led to reduced radial growth in conifers (Pichler and Oberhuber 2007). Another study found increased stem dehydration for conifers compared with broadleaved species, but no detectable change in radial growth with extreme heat (Salomón et al. 2022). This study also found that the timing of the extreme heat event, along with legacy effects of preceding seasons, add complexity to the effects of heatdomes on forests (Salomón et al. 2022). Despite some research into the impacts of heatwaves on forest tree species, the overall impacts of such extreme events on the health of forest ecosystems and the mechanisms that may moderate such effects (like microclimate buffering) remain poorly understood.

To achieve our goal of assessing whether microclimate buffering of forest canopies can minimize the negative impacts of extreme heat events, we addressed three questions.: 1) Does the forest canopy provide consistent buffering during a heatdome, and how does this buffering (during the heatdome) vary with canopy cover? 2) Is buffering great enough to prevent heat stress in forest organisms during the heatdome event? and 3) How do extreme heat events affect canopy cover health?. To address our first question, we compared air and surface temperatures in sites across different levels of canopy cover (from clear-cut to late-successional forests) during the 2021 heatdome in the Pacific Northwest (White et al. 2023). We expected microclimate in late successional forests to be most buffered during the heatdome (less extremes in minima or maxima, and lowest diel variation), and the magnitude of buffering to decrease as canopy cover decreases (Fig. 1). To address our second question, we combined observed temperature data with thermal performance information of 12 species common to the area and assessed whether microclimate buffering was theoretically sufficient to protect these organisms from heat stress, even when temperatures were extremely hot. Finally, we tested the hypothesis that stress following an extreme heat event leads to decreased tree health (e.g., due to heat and water stress) and reduced photosynthetic potential. To answer this, we compared vegetation greenness (Normalized Difference Vegetation Index) following the heatdome in 2021 to the average vegetation index of those same canopies over the same time periods in preceding years.

**Figure 1.**
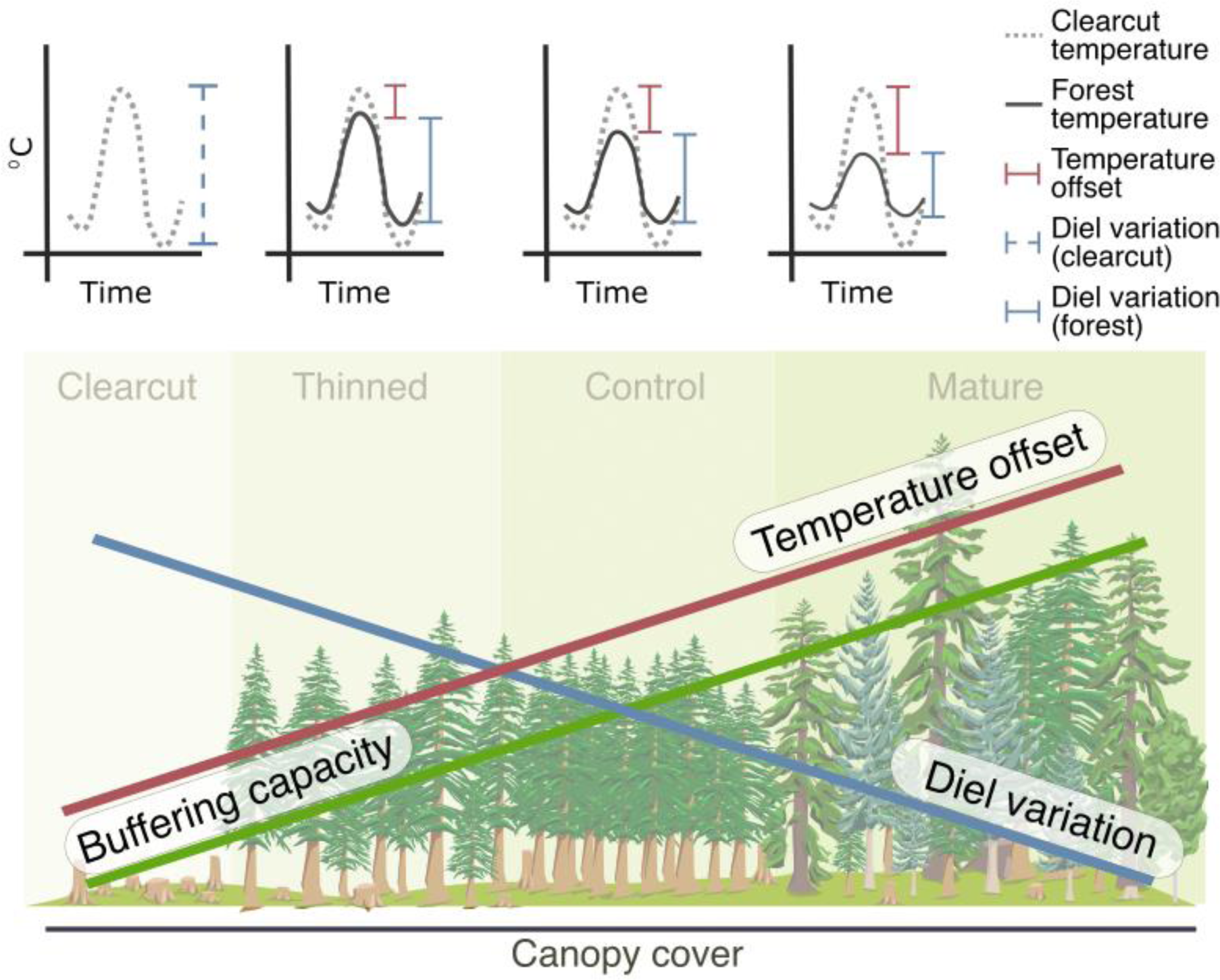
Hypothesized relationships between forest canopy cover and microclimate. As canopy cover and complexity increases from clear-cut to more mature forests, we expect buffering capacity to increase because of which we expect to see decreases in diel variation (blue line) and increases in offset in maximum and mean temperatures (red line). The top panel shows hypothesized differences between temperature in a clear-cut (dashed lines) to temperature in different forest canopy cover.

## Materials and Methods

### Study area

We conducted this study at Ellsworth Creek Preserve (Ellsworth), a 33 km^2^ forested preserve managed by The Nature Conservancy (TNC) located in Willapa Hills (WA, USA). The forests in this region consist of coastal mixed-conifer rainforests that experience warm-dry summers and cool-wet winters. This preserve is part of a watershed-level experiment aimed at restoring former timberlands and accelerating the development of old-growth habitat. As part of this experiment, the Ellsworth Creek watershed was divided into several subbasins with different forest management types such that second-growth forest stands in these sub-basins were either left alone as control (and thus have dense forests and very high canopy cover) or thinned to accelerate development of old-growth forest structure (and thus have relatively lower canopy cover).

### Climate Data

We collected climate data at both micro-(temperature loggers) and macro-(regional climate data) scales. Prior to applying restoration treatments, TNC established permanent vegetation monitoring plots (17.8m radius circular plots; Fig. 2) within all sub-basins. We monitored understory temperatures at a subset of these plots (20 plots; ten in un-thinned (control) and ten in thinned stands) in the North and Central sub-basins (Fig. 2), selected based on topographic position index. Within each plot, we deployed HOBO pendant loggers (Onset Computer Corporation, Bourne, Massachusetts, USA) at two subplots from September 2020 to September 2021 that were programmed to collect temperature data every 2 hours. At each of these two subplots, we monitored surface and air understory temperatures by placing one logger on the ground surface (Ground loggers) and one ∼1.5-2m above the ground (Air loggers). As part of an ongoing study in which we monitored seedling regeneration in a subset of these 20 plots, we deployed two additional ground loggers at six of the 20 plots (Fig 2B). Finally, we also placed loggers in one older (mature) forested stand (two ground and two air) and a clear-cut stand (two ground, no air) to capture the extreme ends of the forest-density spectrum within the preserve.

**Figure 2:**
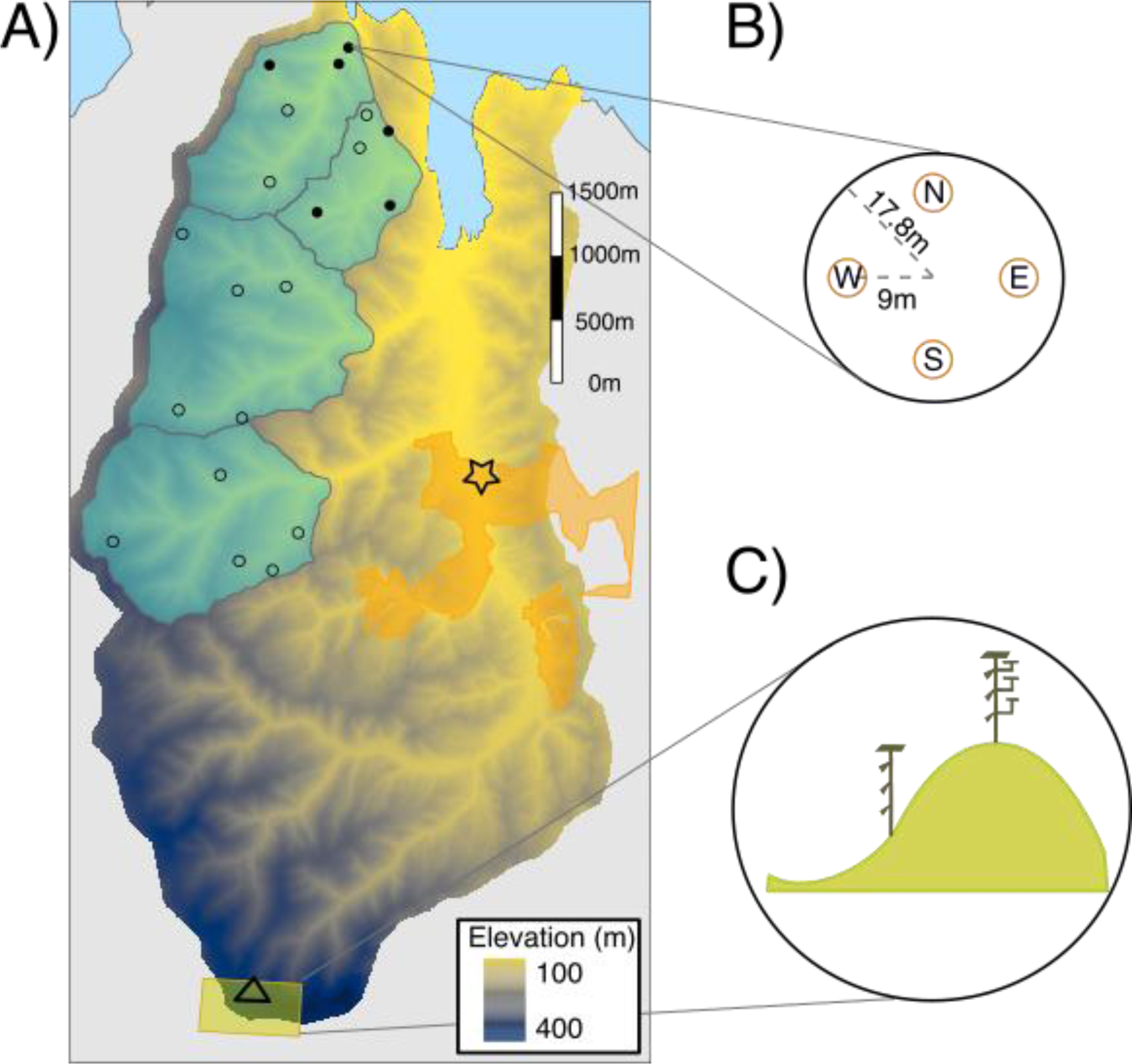
Study design in the forested plots and clear-cut. A) An image of the elevation profile of the study area; and B) the location of circular forested plots (17.8m radius) which were used for collecting ground and air temperatures, B) A close up of forest plots, with orange circles denoting subplots (in each cardinal direction from plot center. C) A close-up of the clear-cut stand (open triangle) which contained two micromet stations - one on the ridge and other one at slope. For all plots used in this study, two subplots were selected (based on canopy cover estimates) for deployment of loggers (A, hollow circles); a ground logger was placed at the selected subplot center and an air logger was tied ∼1.5-2m above the ground. In addition, six plots had additional ground loggers at the remaining two subplots (A, filled circles). The old-growth site is indicated by a star in panel A.

In addition to HOBO pendant loggers, we also established Micromet stations (Dotmote Labs, Seattle, Washington, USA) at two plots (one ridge top, and one at slope; Fig. 2C) in the clear-cut stand. Micromet stations include a solar radiation sensor, wind speed/wind direction and air temperature sensors and are designed for high-spatial temporal collection (every 3 minutes and averaged to bi-hourly intervals; SJ). Wind speed was collected at four different heights (0.3, 0.6, 0.9 and 1.2m) and air temperature was collected at six different heights (0, 0.3, 0.6, 0.9, 1.25, 1.5 m) to capture vertical microclimatic variation. Wind and solar insolation were measured only at the ridge, but temperature profiles were collected at the ridge and slope in the clear-cut.

For our macroclimatic data we obtained regional temperature from ‘gridMet’ (Abatzoglou 2013). This source contains meteorological estimates modeled daily at 4km spatial resolution for the continental United States (CONUS). It combines PRISM and Land Data Assimilation System (NLDAS-2) to provide historic daily reanalysis estimates. To download and process climate estimates, we used the R package *climateR* (https://github.com/mikejohnson51/climateR), which gives daily minimum and maximum temperatures which were averaged to get mean daily temperatures. Fig. 3 shows the mean temperature during the heatdome week across all the landscape types.

**Figure 3.**
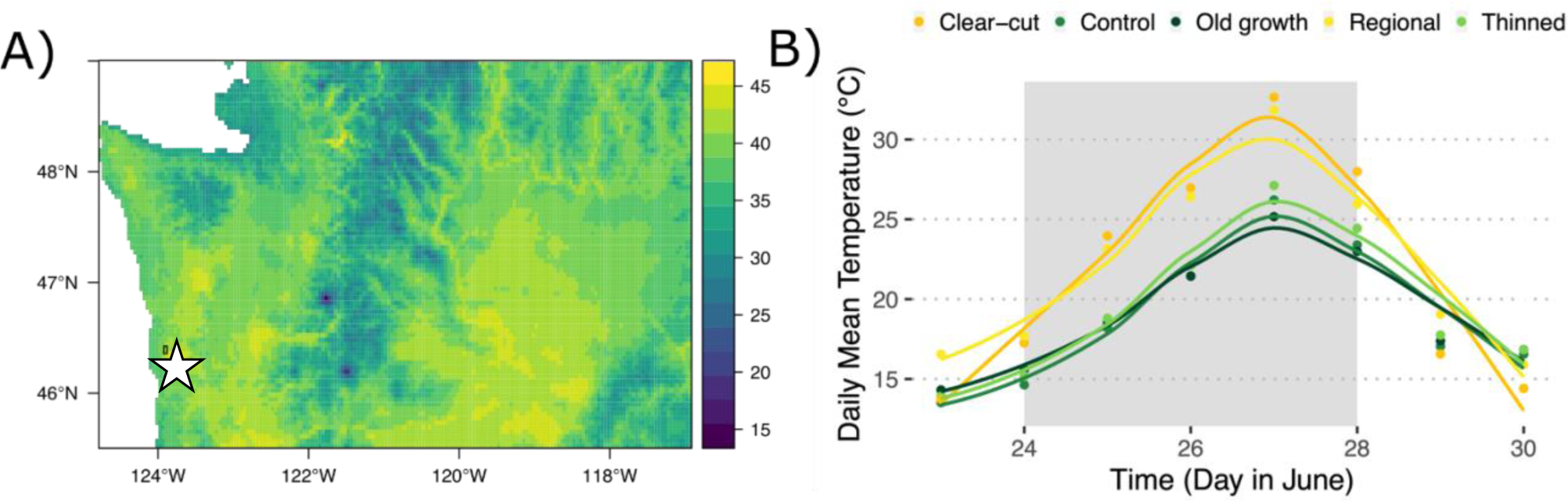
Temperatures during the hottest day (27^th^ June 2021) of the heatdome A) across Washington State, USA (maximum air temperature) from ‘gridMet’, and B) in different forest cover types (Old-growth, Control, Thinned and Clear-cut) at Ellsworth Creek Preserve (boxed area on panel A). In panel B, temperature are mean daily temperatures (averaged across all loggers, both air and soil) in each forest cover type in comparison to regional temperatures (in light yellow). Shaded area in panel (B) refers to heatdome duration.

### Thermal stress data

We used thermal performance data for 12 terrestrial animal species (Supplement B) that occur in the Pacific Northwest (Alaska, Washington, Oregon, Idaho, and British Columbia) to assess how this extreme heat event could impose heat stress on organisms within our forests, and whether such effects may differ depending on canopy cover. First, we collected thermal performance data for 4 species that occur in this region from the *GlobTherm* dataset (Bennett et al. 2018). We also added 8 species listed as species of concern by the Washington Department of Natural Resources and for which we could find Critical Maximum Temperature (CT_max_) data in the literature (Supplement C). We then used CT_max_ of each of these 12 species to compare to distributions of observed temperatures at Ellsworth.

### Remote sensing data

We examined the near-term impacts of the 2021 extreme heat event on forest canopy using remote sensing-based metrics from Planet Labs. Planet Labs (Planet 2018) operates a fleet of satellites called Super Doves which image the earth daily, giving a ground sample distance of 3-5m. PlanetScope, a 4-band level-2 product that gives surface reflectance was downloaded and prepared for analysis using a cloud-agnostic workflow execution provider called SWEEP (John et al. 2019). SWEEP works by running tasks with workflow constructs like scatter, gather, and parallelization. Here we used imagery for 5 years (2017-2021) clipped to a 30m x 30m-pixel area (approx. 900 m square area from the center of the plot) for the 20 sites within both second-growth thinned and control forests. Imagery with more than 15% cloud cover were not included in the analysis. The final dataset from Planet contained information regarding date of collection, site identifier, visible and NIR band statistical metrics (minimum, maximum, mean), and NDVI (minimum, maximum, mean).

### Analysis: Temperature and Buffering Capacity

We calculated daily diel variation, minimum, mean and maximum temperature. We quantified buffering capacity for each of these temperature variables by calculating the temperature offset between understory microclimate (measured by temperature loggers) at our plots, and macroclimate. We used two different metrics of macroclimate: 1) regional gridded temperatures (Abatzoglou 2013) and 2) temperature measured at the micromet stations in the clear-cut site. Microclimate offset can therefore be described by the following equations:

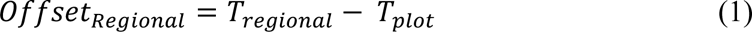

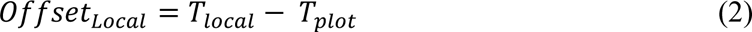

In both the cases, the daily differences between plot temperatures (T_plot_) and either regional temperature (T_regional_) or temperature measured in the clear-cut (T_local_) were used as metrics of buffering capacity such that a higher offset (difference in temperature) means a greater buffering capacity. In other words, a higher offset is equivalent to a cooling effect of the canopy (while a more negative offset indicates a warming effect) when considering minimum, mean, and maximum temperature (equations 1-2). For diel variation, we considered the daily range in temperature (Tmax-Tmin), so that a greater offset indicates a greater reduction in daily variation in temperature within plots.

### Analysis of buffering capacity

We used linear mixed effects models to quantify the buffering capacity of forests with different canopy characteristics during the extreme heat event. We fit models shown in Equations 3 and 4, in which *Offset_Regional_* is a normally distributed response variables of temperature offset for minimum, mean, and maximum daily temperatures and diel temperatures. Explanatory variables were treatment (*x*_1_, consisting of Old-growth, Thinned or Control), age of the stand (*x*_2,_ in years), occurrence of extreme heat event (*x_3_*, Yes or No), and location of the logger (*x*_4_, Air or Ground, only when compared with clear-cut), with interactions between treatment and heat event, and age and heat event. We used plot identity (*u*) as a random intercept and error term is denoted by *e.* The full model specifications are as follows:

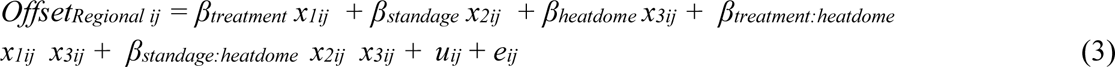

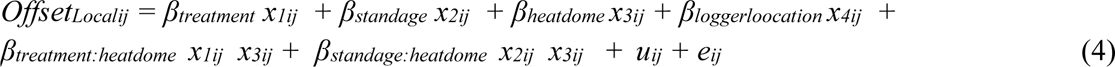

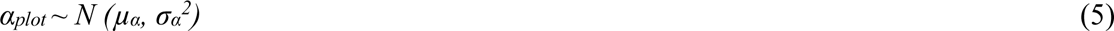

Note that logger location (Air or Ground) was only applicable when we calculated offset relative to temperatures at the clear-cut climate station, because regional gridded climate estimates only provide air temperature. For regional macroclimate comparison, the full month of June was used, and for the local microclimate, partial month of June (June 15-30^th^) was used as data from clear-cut was unavailable for the first half of the month because of technical issues.

### Analysis of species thermal stress

We calculated the total number of hours of heat stress experienced by each of the 12 selected species during the heatdome vs. more average summer conditions. Specifically, we estimated the number of hours observed temperatures exceeded their CT_max_ (Diamond et al. 2012) for a period of two weeks – one immediately before the heatdome (June 17^th^ to 23^rd^) and one during (June 24^th^ to 30^th^). We then calculated the number of heat stress hours (averaged across these 12 taxa) experienced for each of these periods. Additionally, we calculated the percentage of species (of the 12) which experienced any heat stress hour in each of these weeks.

### Analysis of vegetation stress

In coniferous forests, we expect relatively constant NDVI values over a summer without extreme climatic events (Brodrick and Asner 2017, Yang et al. 2019). Canopy level stress, as detected through NDVI, shows up as die-off of vegetation and in coniferous forests is visible as needles turning brown. Therefore, we hypothesized that canopy stress would be apparent as a negative relationship between NDVI and day of the year, indicating a decline in NDVI over the growing season. To detect the impact of the 2021 heatdome on the coniferous forest canopies in our study, we used ordinary least squares regression with mean NDVI obtained from PlanetScope data as the response variable, and day of year (DOY) and treatment type as predictors using the *stats* package (R Core Team 2021). Although more complex, nonlinear relationships between day of year and NDVI would be expected in some systems (e.g., deciduous (Hilker et al. 2014)), we used this approach because it was simple, and a preliminary examination of raw data suggested high variability and an absence of clear seasonal trends in NDVI.

The R statistical environment 4.1.2 (R Core Team 2021) was used for all analyses. For linear mixed effect modeling, ‘lmer’ function in the *lme4* package (Bates et al. 2014), and for data wrangling *tidyverse* package (Wickham et al. 2019) were used.

## Results

### Do forest characteristics influence buffering capacity?

Forested areas—across all forest types--were buffered from hotter temperatures during the heatdome. This was true for comparisons with both clear-cut/local and regional temperatures, and estimated effects of predictors were similar for both comparisons. Thus, we focus on comparison of forest temperatures to clear-cut temperatures (i.e., Offset_Local_) except where relationships diverge between the two comparisons (see Table A.1. for details regarding comparison with Offest_Regional_). The only exception to these differences between microclimate within and outside forests (Offset_Local_ = Temp_local_ – Temp_plot_) demonstrated cooling effects of forests (higher Offset_Local_, meaning greater cooling) (Fig. 4), except for minimum temperatures, which showed no difference (Table 1). Forests of all types were more buffered during the heatdome for all aspects of temperature, except for diel variation when compared to regional temperatures (Table 1 and A.1).

**Figure 4.**
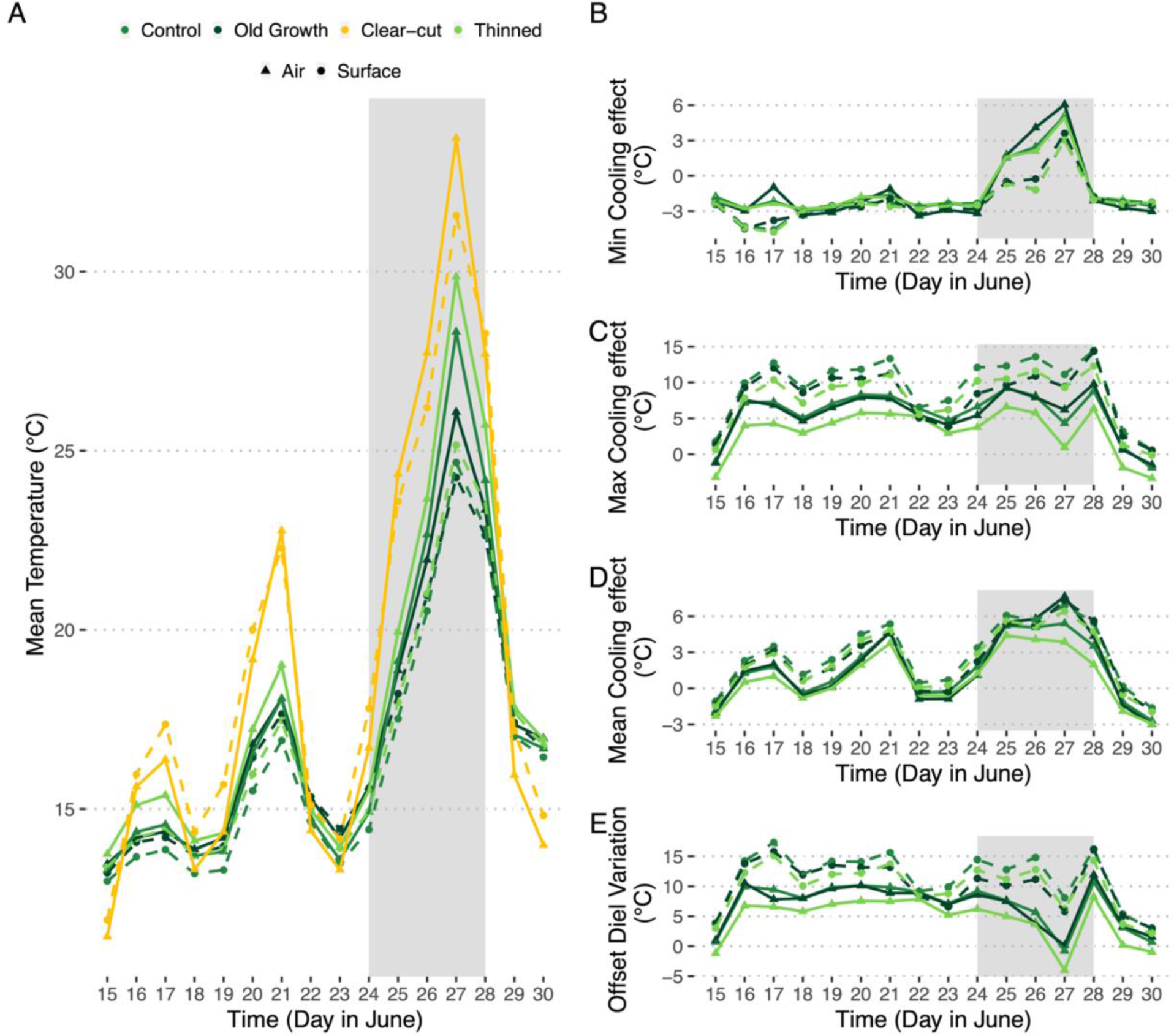
In comparison with the clear-cut, forests exhibit cooling effect and reduced diel variation. Here we show A) observed daily mean temperatures, as well as the difference between temperatures at the clear-cut and within our three kinds of forests (control, thinned, and old-growth) in B) mean, C) min, D) max and E) diel temperature variation. Positive offsets in B-D (offset) demonstrate the cooling effect of forests relative to temperatures at the clear-cut. Positive offsets in E represent a lower diel temperature variation in forests relative to the clear-cut (since offset was calculated as diel variation in clear-cut – diel variation in forest). The shaded areas refer to the heatdome event.

**Table 1:** Summary of linear mixed-effects for the full models fit for local offsets of mean, minima, maxima, and diel variation in temperature. Fixed effects include forest management type (control, thinned, and old-growth), age of the stand, location of logger (ground or air), occurrence of heatdome, and interactions between forest management type and heatdome occurrence, as well as age and heatdome occurrence. Plot was included as an intercept-only random effect. * Indicates significant effects of estimates.

|  | Buffering capacity metrics |  |  |  |
| --- | --- | --- | --- | --- |
|  | Mean Offset (in °C) | Min Offset (in °C) | Max Offset (in °C) | Diel Variation Offset (in °C) |
| Intercept | 0.88*** (0.48, 1.27) | -1.99*** (-2.25, -1.73) | 5.57*** (4.25, 6.89) | 7.58*** (6.21, 8.94) |
| Old Growth | 0.77 (-0.19, 1.74) | 0.01 (-0.61, 0.64) | 2.04 (-1.37, 5.45) | 2.00 (-1.50, 5.49) |
| Thinned | -0.59*** (-0.90, -0.28) | -0.09 (-0.29, 0.11) | -2.06*** (-3.12, -0.99) | -1.95*** (-3.04, -0.85) |
| Age | -0.01*** (-0.02, -0.01) | -0.00 (-0.01, 0.002) | -0.03** (-0.05, -0.01) | -0.02** (-0.05, -0.003) |
| Ground | 1.38*** (1.17, 1.59) | -0.81*** (-0.98, -0.65) | 4.18*** (3.78, 4.59) | 4.96*** (4.49, 5.43) |
| Heatdome | 3.81*** (3.25, 4.37) | 2.42*** (1.98, 2.86) | 4.38*** (3.34, 5.42) | 1.96*** (0.75, 3.17) |
| Old Growth:Heatdome | 0.45 (-0.97, 1.86) | 0.15 (-0.97, 1.27) | 0.59 (-2.03, 3.21) | 0.44 (-2.63, 3.51) |
| Thinned:Heatdome | -0.25 (-0.70, 0.21) | 0.02 (-0.34, 0.38) | -0.38 (-1.22, 0.46) | -0.40 (-1.39, 0.58) |
| Age:Heatdome | 0.00 (-0.01, 0.01) | 0.00 (-0.003, 0.01) | -0.01 (-0.03, 0.01) | -0.01 (-0.03, 0.01) |
| Observations | 1,475 | 1,475 | 1,475 | 1,475 |
| Number of plots | 53 | 53 | 53 | 53 |
| Plot-level std. dev. | 0.30 | 0 | 1.72 | 1.69 |
| Marginal R<sup>2</sup> | 0.47 | 0.40 | 0.33 | 0.25 |

Note: \* p < 0.05, \*\* p < 0.01, \*\*\* p < 0.001 Values in brackets are 95% confidence intervals.

Regarding differences between forest management types, second-growth thinned sites were warmer than second-growth control sites, except for minimum temperatures (treatment was negatively associated with Offset_Local_ magnitude). All forested areas had lower diel variation than clear-cuts with one exception - diel variation in air temperature in the thinned forest was higher than in the clear-cut during the hottest day, although the effect size was not large (Fig. 4E).

Several other predictors also show interesting relationships with Offset_Local_. Surface temperatures were cooler than air temperatures (Table 1); regardless of the treatment status, temperatures were roughly 10 °C cooler on the ground compared to the air on the warmest day (Fig. 4). Minimum temperature Offset_Local,_ however, were similar between the treatments and deviated very little between air and ground. Forest age was negatively associated with Offset_Local_ except for minimum temperatures, i.e., older sites have less of a temperature offset than younger stands.

While the comparisons of forest temperatures with local clear-cut and regional (macro) temperatures were consistent for most of the predictors we assessed, there were differences in some interactive effects. Specifically, forest age and the heatdome had an interactive effect on diel temperature variation when regional temperatures were used for offsetting, suggesting that older forests were less buffered during the heatdome (Table A.1). Similarly, we found an interaction between the thinning treatment and the heatdome event for mean, maximum and diel variations when comparing forest temperature to regional temperature (Table A.1). This suggests that thinned sites were less buffered from the regional macroclimate during the heatdome. By contrast, there was no interactive effect of forest age and the heatdome when examining the difference between forests and local clear-cut temperatures.

### How do canopies insulate species from thermal stress during an extreme heat event?

Although all forest management types were cooler than the clear-cut, the proportion of taxa facing thermal stress (Fig. 5A) varied with canopy cover type, i.e., thinned vs. control forests. Understory organisms experienced similar numbers of hours of thermal stress in both second-growth thinned and second-growth control forests, but a far lower percentage of taxa were predicted to experience stress in the lower canopy in control forests based on air temperatures. However, when we considered surface temperatures there was no difference in percentage of taxa experiencing stress between thinned and control.

**Figure 5.**
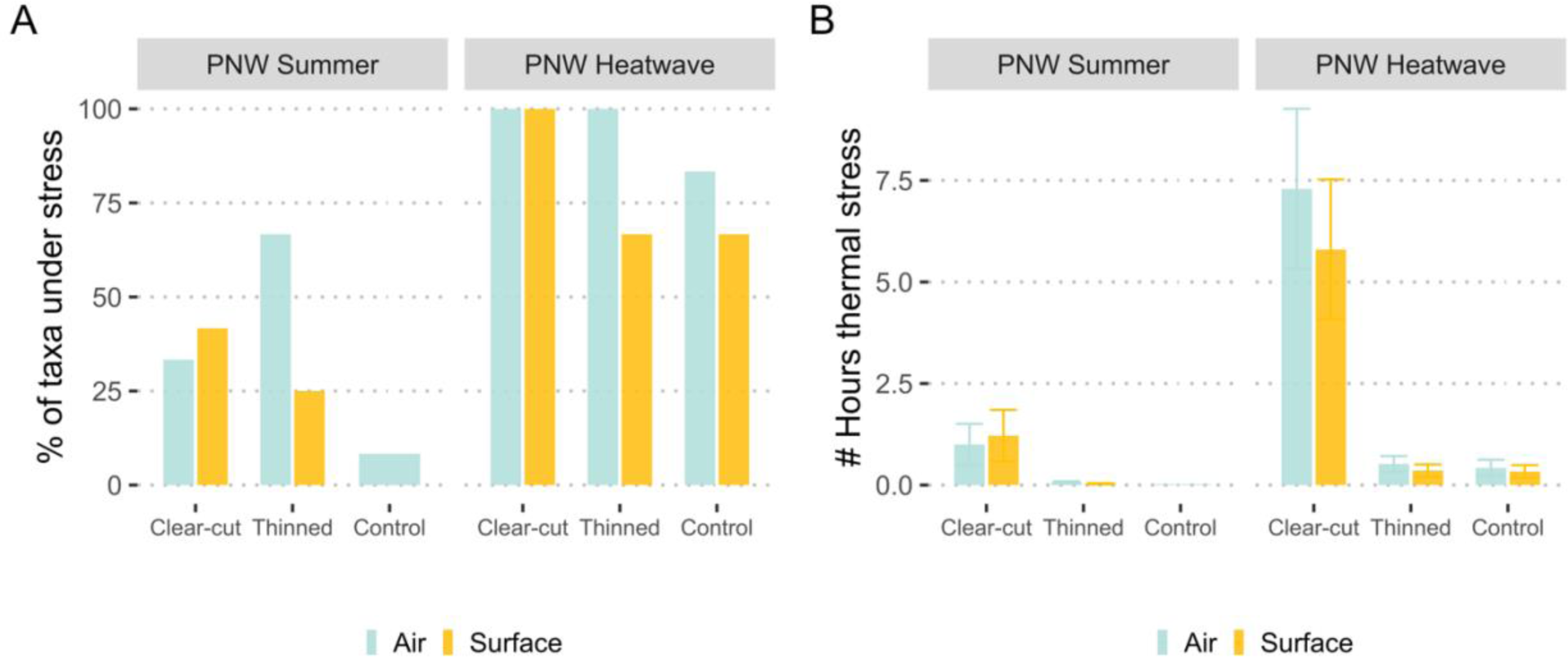
Predicted exposure to thermal stress for 12 common species, before and during the extreme heat event. In A), we show the percent of taxa that were stressed at any point during these two weeks, and in B), the average number of heat stress hours an organism experienced (number of hours a species was subject to temperatures higher than its upper thermal maximum limit).

Predicted thermal stress was greater during the heatdome in all forest types, with nearly all species at risk, and some experiencing many more hours of heat stress per day (e.g., Northern spotted owl (*Strix occidentalis caurina),* Boreal toad *(Bufo boreas) and* Bumble Bee *(Bombus bifarius,* Fig. 5). See Supplement B, especially Fig. B.1, B.2 and B.3, for more details. A clear-cut, which is already expected to be stressful for some terrestrial organisms, becomes even more stressful (approximately double the number of thermal stress hours, Fig. 5B) during the heatdome, while forests are comparatively less stressful (a 66% reduction in hours of thermal stress compared to a clear-cut, and up to 70% fewer taxa experiencing thermal stress: Fig. 5).

### Does forest canopy show signs of stress following an extreme heat event?

Seasonal trends in forest canopy greenness, measured by NDVI, were different during 2021, the heatdome year, compared to other years (Fig. 6). We found that NDVI also varies with canopy cover (Fig. 6B). Specifically, we found that NDVI declined with DOY in 2021 (Pearson’s correlation coefficient: - 0.23; P < 0.01), which contrasts with previous years where there was either no significant relationship between mean NDVI and DOY (2018, Pearson’s correlation coefficient: 0.038, P = 0.17; and 2020, Pearson’s correlation coefficient: 0.027, P = 0.29) or a positive relationship (2017, Pearson’s correlation coefficient: 0.2, P < 0.01; and 2019 Pearson’s correlation coefficient: 0.065, P = 0.014). When grouped by month, the differences in NDVI patterns between 2021 and preceding years is even more apparent (inset Fig. 6A), with post June NDVI exhibiting upward trends in all the years except for 2021.

**Figure 6.**
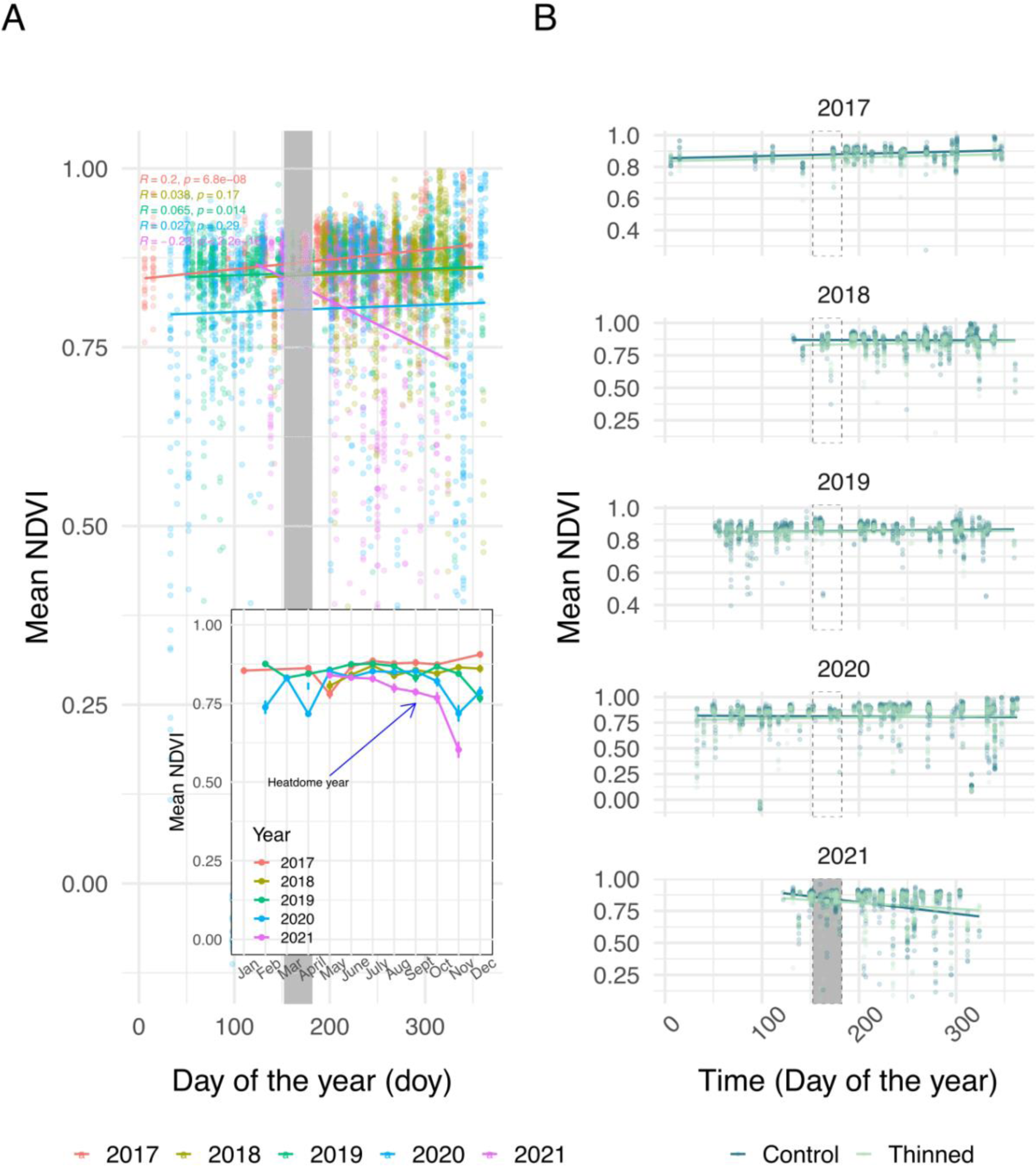
Stress signal in conifers. The dots refer to the plot level NDVI (closer to 1 indicates healthier vegetation). Mean NDVI values regressed against day of the year and inset shows the values by month (A), separated by control, and thinned (light green) vs control (dark green) (B). The smaller number of points in 2017 is because of limited availability of imagery in the early years of Planet product release, and absence of points in first few months of 2018 and in first and last few months of 2021 is because of scene quality and cloud filter. For all years, the grey rectangle refers to June and is shaded in the heatdome year (2021).

## Discussion

Overall, we found that coastal temperate forests buffer understory microclimates during extreme heat events, with forested sites having lower understory temperatures compared to sites without any canopy (from local observations in a clear-cut site) and regional observations (Supplement A). However, we also found that the level of buffering depends on overstory characteristics, indicating that heterogeneity in forest canopy characteristics (in our case, canopy cover) also creates heterogeneity in understory microclimate, similar to findings from a previous study on stand structure characteristics (Gril et al. 2023), and aligns with our expectations that greater canopy cover leads to greater buffering of understory temperatures. Our results suggest that these moderated understory temperatures could lead to lower heat-stress for understory organisms experiencing a heatdome. Finally, we found that extreme heat events also stressed forest canopies during the heat dome year. We discuss these findings as well as their implications for conservation and management in the following sections.

### Forest characteristics influence buffering capacity

Buffering effects of forests were more pronounced during the heatdome event for all aspects of temperature (positive coefficients for Offset_Local_; Table 1). Thus, in addition to decoupling the understory climate from the broader macroclimate during the relatively mild summer temperatures experienced in the Pacific Northwest (Davis et al. 2018b), we found forests maintain their buffering capacity even during extreme heat events, where temperatures were >16°C warmer than normal (White et al. 2023). Specifically, in our study forests were, on average, 3°C cooler than a clear-cut area and 4°C cooler than regional temperatures during the heatdome (Fig. 4). These temperature offsets are comparable those from a global analysis of paired within and outside forest temperature loggers (De Frenne et al. 2019) with a key difference being that the average cooling effect found at our study sites during a heatdome were closer to the cooling effect on maximum average temperatures. However, there was variation in temperature related to canopy cover, which is lower in second-growth thinned forests compared to second-growth control sites, resulting in a higher amount of radiation reaching the forest understory (Oke 1976). This presumably explains the elevated temperatures we quantified during the heatdome event, and similar to what occurs during warmer months (Zellweger et al. 2019). The cooling effect we observed in managed forests was comparable to the cooling effect we observed within our old-growth stand, particularly in stands of the second-growth forests where the canopy had not been thinned (controls), demonstrating the key role of canopy cover in determining how much buffering is possible (Chamberlain et al. 2021, Au et al. 2022). One additional point to note is that the during heatdome, microclimate in the forest understory can be influenced by convection. Convection, driven by temperature differences, has the capacity to shape microclimates by creating convection currents that transport heat away from near the surface (MacHattie and McCormack 1961, Oke 1976). Overall, this suggests that forests can buffer the understory microclimate even from extreme heat events.

In addition to horizontal heterogeneity, we also found that microclimate buffering varied vertically, with ground temperatures generally cooler than air temperatures by ∼10°C (Fig. 4 and Supplement A). This contrasts with vertical temperature patterns in the clear-cut, where surface temperatures gradually exceed air temperature as the day progresses (Oke 1976). Our findings suggest an inversion-like pattern (temperature increase with height) during mid-day (Jarvis 1976, Oke 1976, De Frenne et al. 2021) which increases at a faster rate in second-growth thinned sites compared to control sites. More broadly, this implies that the buffering capacity of forests, whether old-growth or thinned, are more likely to moderate heatdome effects for ground-dwelling organisms (e.g., ground beetles, amphibians, seedlings) than for those operating at mid- and upper-canopy levels (e.g., birds).

Surprisingly, younger stands had greater temperature offsets (i.e., were cooler than clearcuts or regional temperatures) than our old-growth site, seeming to contradict our assumption that old-growth or late successional stands provide more buffering ((Lindenmayer et al. 2022); Fig. 3, Fig. 4, and Fig. A.1). One potential reason for this finding is that the canopy complexity (which we assume buffers understory temperatures) develops over time differently in these managed forests than the more “natural” old-growth forests in which the development of canopy complexity has been studied (Chamberlain et al. 2021), and one old-growth site that was used for reference. Typical of intensively managed forests, Ellsworth consists of forests that were planted to be dense while still permitting radial tree growth for efficient timber production, leading to a relatively high canopy overlap in younger forests – likely much higher than during unmanaged succession in these forests. A second point is that our finding that the magnitude of offsets depended on whether we compared forest temperatures to local clearcut temperatures or regional temperatures. This illustrates a key point - the reference used for comparison with forest understory climate matters. Thus, it is important to consider which temperatures to use as a reference when exploring microclimate buffering (Maclean et al. 2021, Haesen et al. 2021, Gril et al. 2023), with differing results possible if choosing a far-away weather station vs. a gridded climate product vs. a local station that is embedded within the study area.

### Microclimate buffering can moderate thermal stress during extreme heat

Our findings show that forests can act as a refuge from extreme heat for forest organisms, highlighting the importance of managing and restoring forests and maintaining canopy cover to minimize the negative impacts that extreme heat events can have on organisms, communities, and ecosystems (Harris et al. 2018, Maxwell et al. 2019). Even during an average PNW summer, we found that understory organisms would face some degree of thermal stress, as demonstrated by the average number of hours of thermal stress experienced and the proportion of taxa facing thermal stress (Fig. 5A, B). We found that extreme heat events may push the limits of organismal thermal tolerance, but high canopy cover forests offer the possibility of moderation by canopy cover. The degree of heat stress and number of organisms experiencing heat stress also varied depending across surface versus air temperature. This highlights the importance of examining vertical heterogeneity in buffering (Haesen et al. 2021), since the microclimate that an organism experiences depends on what part of vertical space it inhabits.

Future studies could allow for an even more nuanced exploration into these dynamics, potentially aiding in our understanding of which species might be particularly affected by extreme heat events. For instance, the presence of vertical and horizontal heterogeneity in buffering within forests suggests that impacts will depend on not only organismal thermal limits, but also the parts of the forest they occupy. Similarly, the duration of extreme heat events may also matter, as survival has been shown to depend on the duration of a heat stress in addition to the intensity (Rank et al. 2022). Over longer time periods, while increases in incidence and magnitude of episodic extreme heat events could influence changes in thermal tolerances of some organisms (Sunday et al. 2019), it would be crucial to distinguish between the taxa that might be able to adapt vs. those that might not. In addition, organisms better able to behaviorally respond to heatdome events by seeking out cooler temperatures, either due to movement capability or behavioral adaptations, may also be better able to take advantage of canopy buffering.

### Forest canopy is under stress following extreme heat

NDVI has been useful in detecting signals of canopy stress following disturbances and other extreme events (like soil attrition (Zhang et al. 2020), water deficit (Lobo and Maisongrande 2006), and insect damage (Foster et al. 2017). In recent years, NDVI has also been used to examine the effect of extreme heat events on forest canopy (Pettorelli et al. 2005, Yuhas and Scuderi 2009). The declining NDVI responses we observed in our sites following the heat dome event (Fig. 6) are similar to NDVI responses to longer term droughts observed in previous studies (Yoshida et al. 2015, Song et al. 2019, Pompa-García et al. 2021). Thus, our results suggest that canopy stress occurs in response to the extreme climatic conditions of a heatdome (as measured by NDVI). However, we also found that the magnitude of canopy stress varies across treatment types, with control sites showing a greater decrease in NDVI compared to thinned sites in heatdome years (although the difference was not large – Fig. 6). These results could indicate that forests with greater tree density are more vulnerable to heat-stress during extreme heat events. However, NDVI values depend on other factors which may be responsible for this result – like the radiometric calibration of the sensor, and environmental forcing like precipitation (Santos and Negri 1997). Coastal coniferous forests like the ones in our study are in a high precipitation zone and have a long growing season; declining NDVI in denser canopies following the heatdome might have been complicated by saturation in optical signal caused by shading induced by denser canopies (Yoder and Waring 1994, Leblanc et al. 1997, Brodrick and Asner 2017).

In drought-impacted or semi-arid regions, increases in canopy temperatures have resulted in adult tree mortality (Scherrer et al. 2011, Guha et al. 2018), but it is unclear whether heat-induced tree mortality will occur in these coastal temperate forests following heatdomes, because such events have been extremely rare in the Pacific Northwest. However, extreme heat events are expected to increase in frequency in the Pacific Northwest with climate change, so the potential impacts of increased canopy stress on tree mortality following individual or multiple heatdome events needs to be monitored. These episodic events will likely exacerbate the effects of long-term warming (and *vice* versa) through press and pulse dynamics (Harris et al. 2018) and have potential to cause damage beyond the effects of a single extreme event alone. Moreover, less dramatic effects on forests, including crown die-back following heatdome events, can also alter the forest’s buffering capacity (Ruthrof et al. 2016). More broadly, if the increased plant stress (as indicated by the decrease in NDVI, Fig. 6) we observed after the 2021 heatdome event eventually deteriorates the ability of forests to buffer the understory from extreme heat, this could spell trouble for understory organisms already coping with the negative impacts of climate change.

## Conclusions

As the frequency and duration of episodic extreme heat events increase (IPCC 2022), the persistence of forest understory microclimatic conditions that can create critical refugia in a warming climate may be tested. Our study quantifies microclimate buffering capacity of forests during the 2021 PNW heatdome, and we demonstrate that forests can buffer understory temperatures during extreme heat events. Specifically, temperature differences between forested and unforested sites were substantial enough that the heat stress experienced by terrestrial organisms during the heat dome would be significantly moderated by forest cover. However, our study also highlights that not all forests have similar buffering capacities. Understories of thinned second-growth forests experienced a greater increase in temperature (∼ 5°C warmer and more variable) than those of un-thinned second-growth forests during the heatdome of 2021. More worryingly, we also detected vegetation stress in response to the heat dome, implying that the ability of forests to buffer understory temperatures from extreme heat dome events may be threatened by repeated exposure to extreme heat.

On a more practical level, we believe that a better understanding of the relationship between forest cover and buffering during extreme heat events could provide insight into best practices for forest restoration and management in a world characterized by higher temperatures and more frequent extreme events. For example, the canopy cover in our study system was manipulated as part of an experiment to understand how and whether thinning could accelerate the restoration of old-growth forest structure, and our results suggest that removing canopy cover has negative impacts on organisms experiencing heatwaves. Thus, it is possible that as these heatdome events increase in frequency and magnitude given warming trends (Guerreiro et al. 2018, Clarke et al. 2022), their increased negative impacts on thinned forests (e.g. through impacts on understory organisms of interest and / or potentially forest regeneration) could interfere with conservation and restoration goals. This result may seem contradictory to that of a related study (Pradhan et al. 2023) examining climate change refugia (areas of the landscape relatively buffered from climate change impacts (Morelli et al. 2016)), where management through thinning did not alter the buffering capacity in the Ellsworth Creek watershed. The differing conclusions reached by examining the results of these two studies could arise due to different methods (e.g., spatio-temporal scale, use of proxy measures vs. temperature). However, we think it is more likely that there are differences in how canopy thinning influences extreme temperatures (explored in this study) vs average seasonal temperatures (Pradhan et al. 2023). Collectively, these two papers therefore highlight the complexity of decision-making for conservation and the importance of considering specific conservation goals. One valuable approach may be to modify traditional density reduction treatments with knowledge of how organisms of interest experience extreme thermal heat stress in different parts of the forest. If this is possible, managers may be better able to balance restoration goals achieved through density reduction while maintaining resilience to climate change.

## Author contributions

J.H.R.L, K.P., M.C., A.K.E. and A.J. designed the study and collected the data. A.J. and K.P. analyzed the data with assistance from A.K.E. and J.H.R.L. A.J. and K.P. developed the first draft of the manuscript. All the authors contributed to reviewing and editing the draft.

## Funding

This study was funded by grants from Department of Biology, University of Washington to A. J., a U.S Geological Survey Northwest Climate Adaptation Science Center award G17AC00218 to K.P., a University of Washington Biology Department graduate fellowship to K.P., a U.S. National Science Foundation Award to J.H.R.L and Amy L. Angert (DEB-1555883). We also would like to thank ETH Zurich for support during development of this manuscript.

## Declaration of Competing Interest

The authors declare that they have no competing interests.

## Acknowledgments

We are grateful to the Planet Labs and Microsoft AI for Earth program for providing the satellite imagery and computational support. We also thank the TNC for the opportunity to conduct world-class climate research in their sustainable forests. We acknowledge that some of this research was conducted in ancient lands of Lower Chinook people, the land which touches the shared waters of all tribes and bands within the Confederated Tribes of Grand Ronde, Chinook, and Confederated Tribes of Siletz Indians.

## Notes

### Competing Interest Statement

The authors have declared no competing interest.

