## Supplement A for "Microclimate buffering varies across forest types during an extreme heat event"

#### Buffering capacity of forests in relation to regional macroclimate

Forest temperatures are cooler than regional temperatures during the extreme heat event (Fig. A.1). Table A.1 shows the full model for the month of June including the week of heatdome (June 24-28$^{th}$). Old-growth sites have higher $Offset_{Regional}$ than control sites suggesting old-growth forests are more buffered than control sites while thinned sites have a smaller $Offset_{Regional}$ than control sites for all aspects of temperature other than minimum temperatures (Fig. A.1). Stand age is negatively associated with $Offset_{Regional}$ for mean and max temperature, and diel variation, but did not strongly affect offset in minimum temperatures. The offsetting of mean, minimum and maximum temperatures within forests was higher during the heatdome (>2 °C cooler) but did not influence offset in diel variation. The interaction between treatment and heatdome status affected the mean, max and diel temperatures, but not the minimum temperatures. Thinned and heatdome interaction affected mean, maximum, and diel temperatures negatively in comparison to control sites, whereas interaction between old-growth and heatdome was non-significant.  Stand age and heatdome interaction was significant only for diel variation.

Table A.1: Linear mixed-effects for the full model for regional offsets of mean, minima, maxima and diel variation in temperatures. Fixed effects include forest management type (control, thinned, and old-growth), age of the stand, occurrence of heatdome, and interactions between forest management type and heatdome occurrence as well as age and heatdome occurrence. Plot was included as an intercept-only random effect. $^{*}$ Indicates significant effects of estimates.

| | Buffering capacity metrics | | | |
|---|---|---|---|---|
| | Mean Offset (in °C) | Min Offset (in °C) | Max Offset (in °C) | Diel Variation Offset (in °C) |
| Intercept | 1.29$^{***}$ (0.90, 1.68) | -2.90$^{***}$ (-3.21, -2.60) | 5.45$^{***}$ (4.06, 6.84) | 8.29$^{***}$ (6.85, 9.73) |
| Old Growth | 0.94$^{**}$ (0.01, 1.88) | -0.07 (-0.79, 0.66) | 2.98$^{*}$ (-0.41, 6.37) | 3.03$^{*}$ (-0.48, 6.53) |
| Thinned | -0.59$^{***}$ (-0.91, -0.28) | 0.03 (-0.22, 0.27) | -2.23$^{***}$ (-3.36, -1.11) | -2.22$^{***}$ (-3.39, -1.05) |
| Age | -0.01$^{***}$ (-0.02, -0.004) | 0.00 (-0.004, 0.01) | -0.03$^{***}$ (-0.06, -0.01) | -0.03$^{***}$ (-0.06, -0.01) |
| Heatdome | 2.45$^{***}$ (1.87, 3.03) | 2.62$^{***}$ (1.88, 3.36) | 3.19$^{***}$ (2.41, 3.96) | 0.57 (-0.42, 1.56) |

| | | | | |
|---|---|---|---|---|
| Old Growth:Heatdome | 0.81 (-0.58, 2.20) | -0.06 (-1.85, 1.72) | 1.36 (-0.51, 3.23) | 1.42 (-0.95, 3.80) |
| Thinned:Heatdome | -0.60** (-1.06, -0.13) | -0.03 (-0.63, 0.57) | -0.83*** (-1.46, -0.21) | -0.81** (-1.60, -0.01) |
| Age:Heatdome | -0.00 (-0.01, 0.01) | 0.00 (-0.01, 0.02) | -0.01 (-0.02, 0.004) | -0.01* (-0.03, 0.002) |
| Observations | 1,173 | 1,173 | 1,173 | 1,173 |
| Number of plots | 40 | 40 | 40 | 40 |
| Plot-level variance | 0.38 | 0 | 1.69 | 1.73 |
| Marginal $R^2$ | 0.27 | 0.24 | 0.27 | 0.16 |
| Note: | | | | *p**p***p<0.01 |
| | | | | Values in brackets refer to 95% confidence intervals. |

Figure A.1. Here we show A) daily mean temperatures in the region, forests (control) and thinned forests, as well as the difference between regional temperatures and our two kinds of forests (control and thinned) in B) mean, C) min, D) max and E) diel temperature variation. Positive offsets in B-D (offset) demonstrate the cooling effect of forests relative to regional temperatures; and positive offsets in E represents a lower diel temperature variability in forests relative to the regional diel variability. The shaded area is the heatdome event.

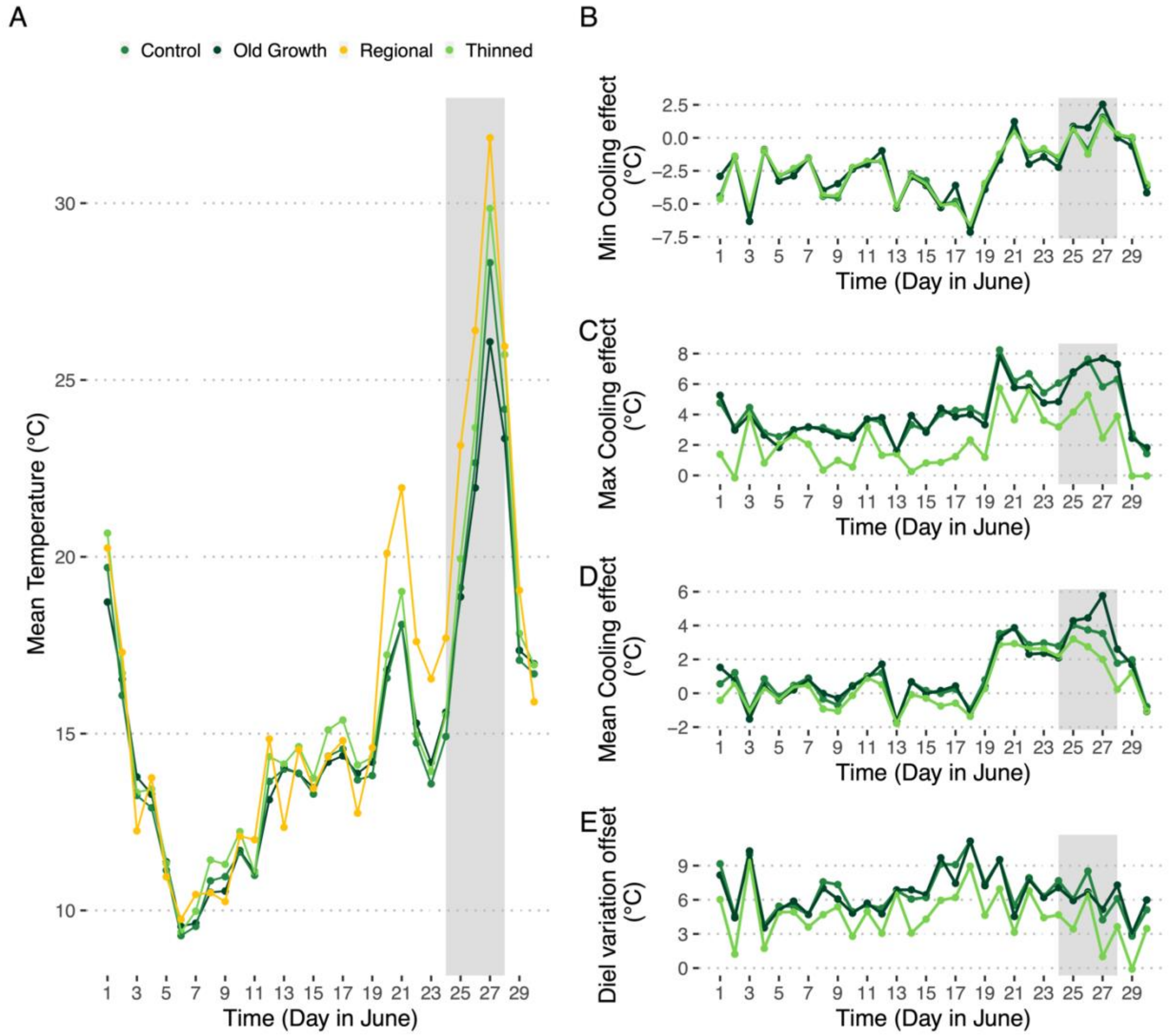


### Supplement B

#### Thermal stress during extreme heat at species level

Understory organisms likely face a thermal stress even during an average Pacific Northwest summer, as demonstrated by the average number of hours of thermal stress faced and the proportion of taxa facing thermal stress across the three landscape types.

Figure B.1. Species exposed to stress events in the clear-cut; each dot refers to the number of hours in the study area (clear-cut specifically) the species was subject to temperatures higher than its upper thermal max limit. The column headers refer to the day in June 2021, and row headers signify either Air or Surface temperatures. The size of the dot does is informing the reader of the higher stress events reported on the hottest day.

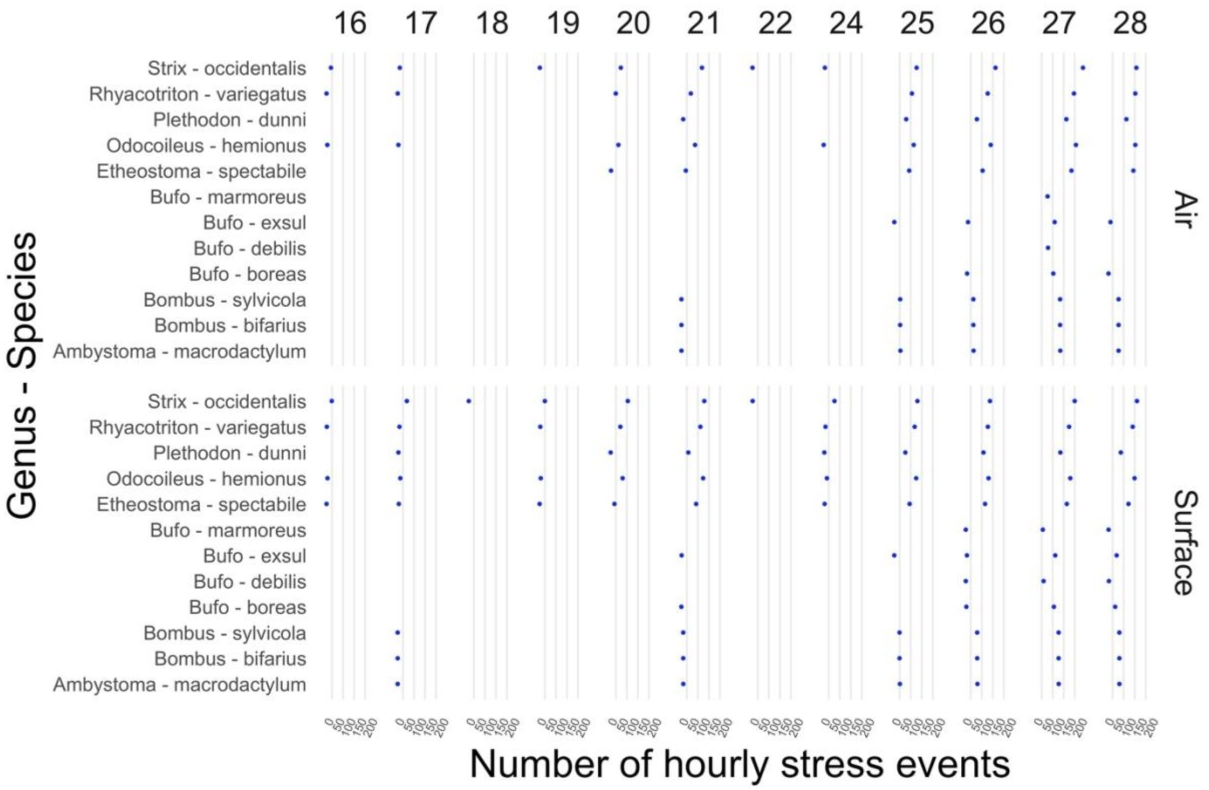


Figure B.2. Species exposed to temperature stress events in the Thinned treatment; each dot refers to the number of hours in the study area (clear-cut specifically) the species was subject to temperatures higher than its upper thermal max limit. The column headers refer to the day in June 2021, and row headers signify either Air or Surface temperatures.

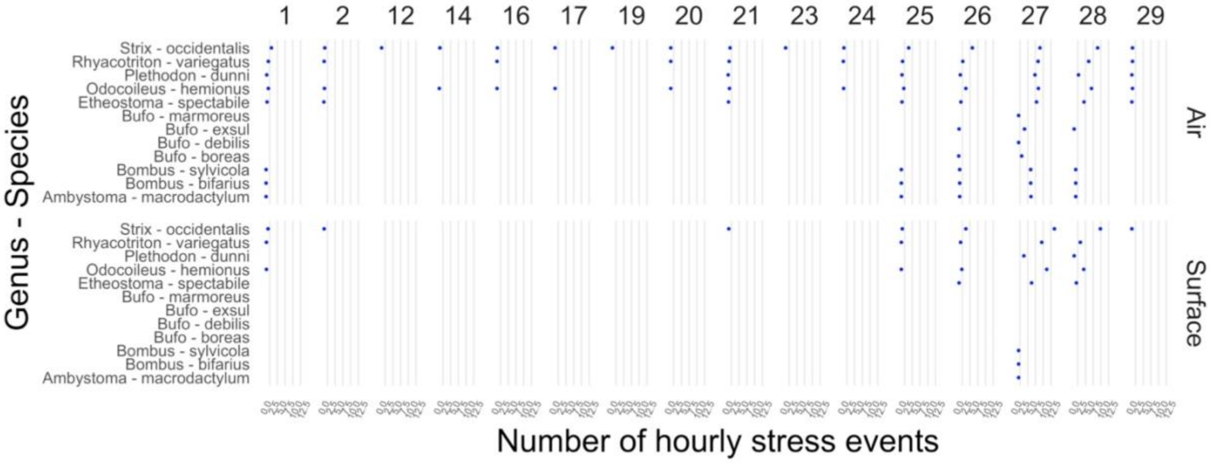


Figure B.3. Species exposed to temperature stress events in the Control treatment; each dot refers to the number of hours in the study area (clear-cut specifically) the species was subject to temperatures higher than its upper thermal max limit. The column headers refer to the day in June 2021, and row headers signify either Air or Surface temperatures.

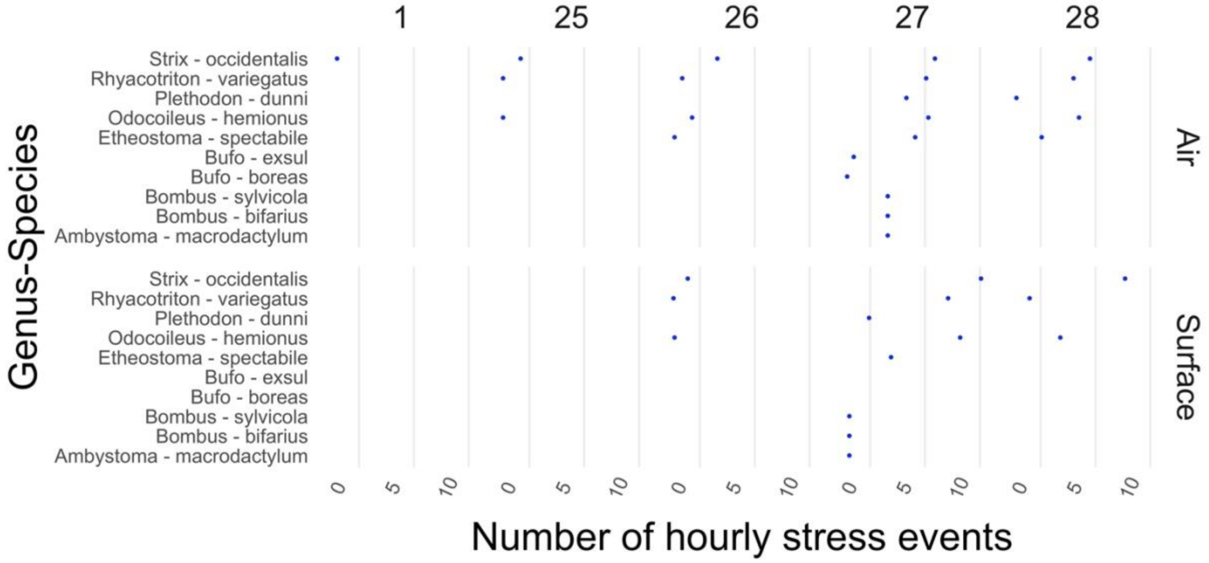
