## Supplement B for "Microclimate buffering varies across forest types during an extreme heat event"

### Thermal stress during extreme heat at species level

Understory organisms likely face a thermal stress even during an average Pacific Northwest summer, as demonstrated by the average number of hours of thermal stress faced and the proportion of taxa facing thermal stress across the three landscape types.

Figure B.1. Species exposed to stress events in the clear-cut; each dot refers to the number of hours in the study area (clear-cut specifically) the species was subject to temperatures higher than its upper thermal max limit. The column headers refer to the day in June 2021, and row headers signify either Air or Surface temperatures. The size of the dot does is informing the reader of the higher stress events reported on the hottest day.

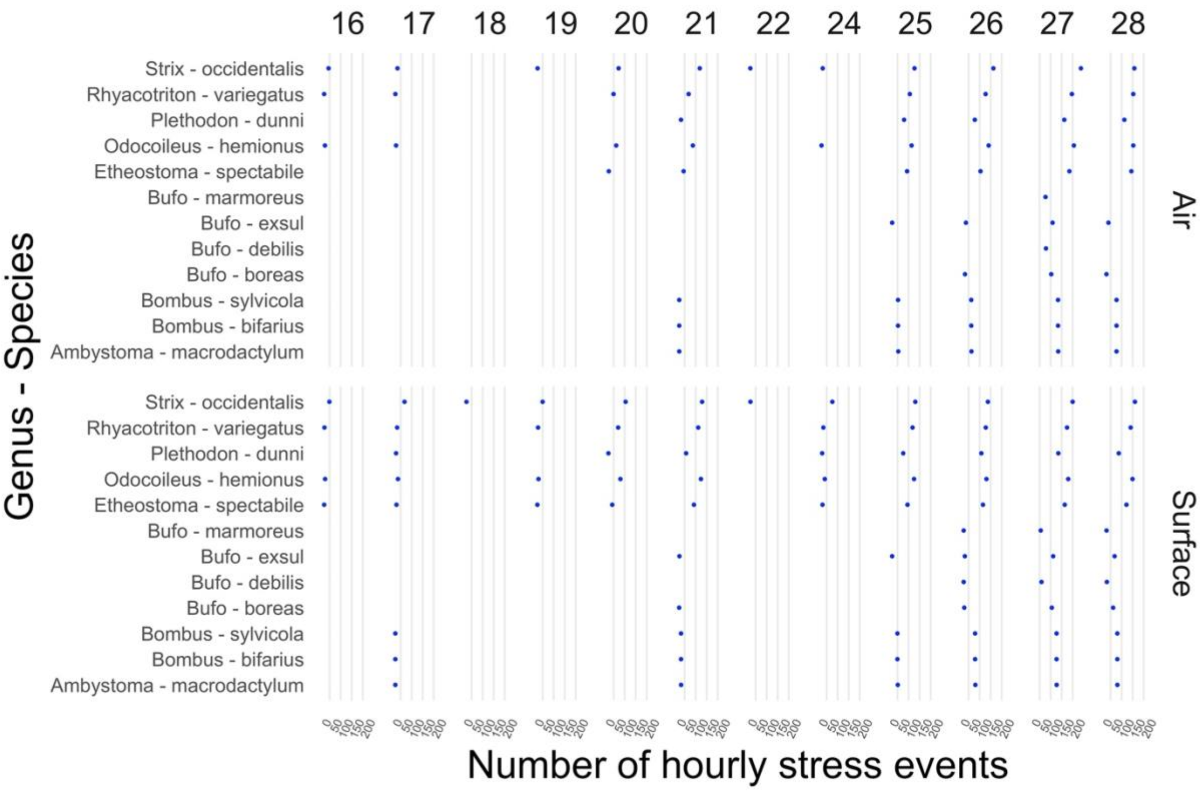


Figure B.2. Species exposed to temperature stress events in the Thinned treatment; each dot refers to the number of hours in the study area (clear-cut specifically) the species was subject to temperatures higher than its upper thermal max limit. The column headers refer to the day in June 2021, and row headers signify either Air or Surface temperatures.

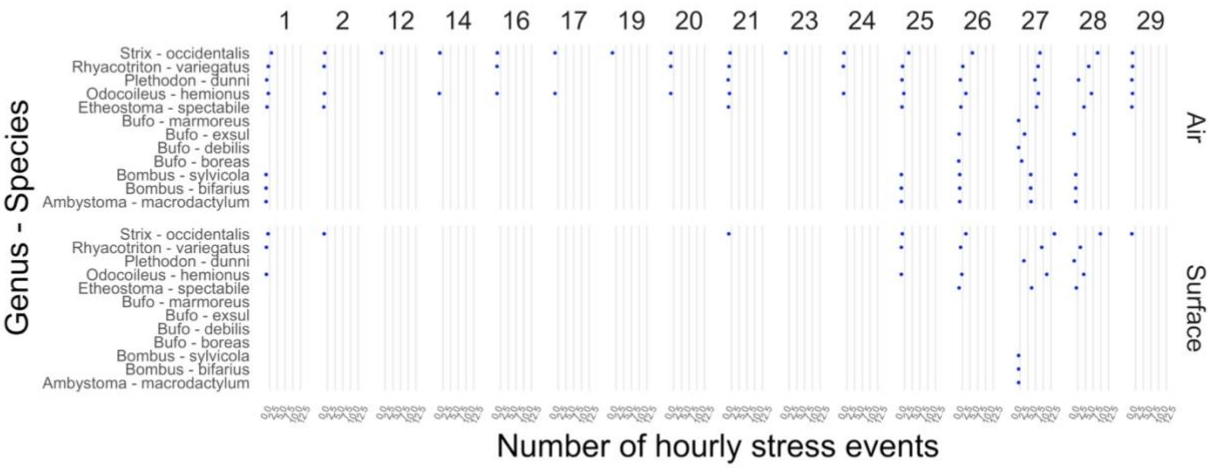


Figure B.3. Species exposed to temperature stress events in the Control treatment; each dot refers to the number of hours in the study area (clear-cut specifically) the species was subject to temperatures higher than its upper thermal max limit. The column headers refer to the day in June 2021, and row headers signify either Air or Surface temperatures.

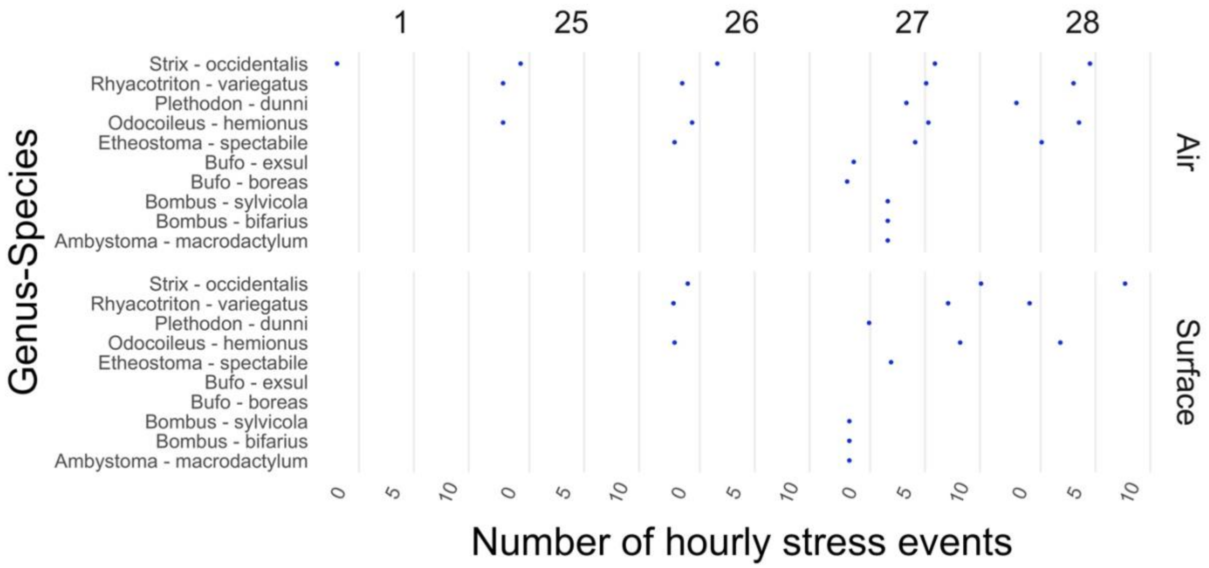
